# Comparative and population genomics landscape of *Phellinus noxius*: a hypervariable fungus causing root rot in trees

**DOI:** 10.1101/132712

**Authors:** Chia-Lin Chung, Tracy J. Lee, Mitsuteru Akiba, Hsin-Han Lee, Tzu-Hao Kuo, Dang Liu, Huei-Mien Ke, Toshiro Yokoi, Marylette B Roa, Meiyeh J Lu, Ya-Yun Chang, Pao-Jen Ann, Jyh-Nong Tsai, Chien-Yu Chen, Shean-Shong Tzean, Yuko Ota, Tsutomu Hattori, Norio Sahashi, Ruey-Fen Liou, Taisei Kikuchi, Isheng J Tsai

## Abstract

The order Hymenochaetales of white rot fungi contain some of the most aggressive wood decayers causing tree deaths around the world. Despite their ecological importance and the impact of diseases they cause, little is known about the evolution and transmission patterns of these pathogens. Here, we sequenced and undertook comparative genomics analyses of Hymenochaetales genomes using brown root rot fungus *Phellinus noxius*, wood-decomposing fungus *Phellinus lamaensis*, laminated root rot fungus *Phellinus sulphurascens*, and trunk pathogen *Porodaedalea pini*. Many gene families of lignin-degrading enzymes were identified from these fungi, reflecting their ability as white rot fungi. Comparing against distant fungi highlighted the expansion of 1,3-beta-glucan synthases in *P. noxius*, which may account for its fast-growing attribute. We identified 13 linkage groups conserved within Agaricomycetes, suggesting the evolution of stable karyotypes. We determined that *P. noxius* has a bipolar heterothallic mating system, with unusual highly expanded ~60 kb *A* locus as a result of accumulating gene transposition. We investigated the population genomics of 60 *P. noxius* isolates across multiple islands of the Asia Pacific region. Whole-genome sequencing showed this multinucleate species contains abundant poly-allelic single-nucleotide-polymorphisms (SNPs) with atypical allele frequencies. Different patterns of intra-isolate polymorphism reflect mono-/heterokaryotic states which are both prevalent in nature. We have shown two genetically separated lineages with one spanning across many islands despite the geographical barriers. Both populations possess extraordinary genetic diversity and show contrasting evolutionary scenarios. These results provide a framework to further investigate the genetic basis underlying the fitness and virulence of white rot fungi.

## Introduction

Under most circumstances, fungi coexist with trees or act as saprotrophs responsible for carbon and nitrogen cycling in forest systems. However, some fungi are also one of the dominant groups of pathogens causing diseases in trees. There has been an emergence of tree disease outbreaks in different parts of the world such as ash dieback (Gross, Holdenrieder, Pautasso, Queloz, & Sieber, 2014), Dutch elm disease (Potter, Harwood, Knight, & Tomlinson, 2011), laminated root rot caused by *Phellinus sulphurascens* (Williams et al., 2014), and brown root rot caused by *Phellinus noxius* (Akiba et al., 2015; P. Ann, Chang, & Ko, 2002; Chung et al., 2015). Factors contributing to this phenomenon include climate change (Goberville et al., 2016) and human activity (Fisher et al., 2012). If interventions are not implemented early and effective, pathogen infections can kill millions of trees and the spread can become very difficult to stop (Cunniffe, Cobb, Meentemeyer, Rizzo, & Gilligan, 2016).

Hymenochaetales is dominated by wood decay fungi and belongs to Agaricomycetes of Basidiomycota. Most species within this order are saprotrophic but some also exhibit pathogenic lifestyles that have been recorded in major forest incidents as early as 1971 in different parts of the world (Hepting, 1971; Norio Sahashi, Akiba, Ishihara, Ota, & Kanzaki, 2012). In particular, *P. noxius* has a very wide host range, spanning more than 200 broadleaved and coniferous tree species (at least 59 families) including many agricultural, ornamental, and forest trees such as longan, litchi, camphor, banyan, and pine (P. Ann et al., 2002; Norio Sahashi et al., 2014). Inoculation assays showed that only seven out of 101 tested tree cultivars (92 species) exhibited high resistance (P. J. Ann, Lee, & Huang 1999). Despite the recognized importance of *P. noxius* as an emerging pathogen, its genome, evolution and global population genetics is poorly understood. Previous reports based on simple sequence repeat (SSR) markers suggest the existence of highly diversified *P. noxius* populations (Akiba et al., 2015; Chung et al., 2015), but the isolates exhibited little to no host specificity (P. J. Ann et al., 1999; Nandris, Nicole, & Geiger, 1987; N. Sahashi, Akiba, Ishihara, Miyazaki, & Kanzaki, 2010). Currently, no gold standard genomes (Chain et al., 2009) are available for this group of wood decay fungi, which is a necessary step toward a better knowledge of their diversity, ecology and evolution.

The life cycle of *P. noxius* (P. Ann et al., 2002) is thought to be similar to other important root-rotting basidiomycetes, such as *Armillaria mellea* and *Heterobasidion annosum*. The new infection can start from previously infected plants/stumps or colonized wood debris, from which the mycelium of *P. noxius* grows to infect the lateral and tap roots of the host tree (P. Ann et al., 2002). An invasion to the cortex and lignified xylem is usually observed (T.-T. Chang, 1992), sometimes accompanied by gradual expansion of the mycelial mat to the basal stem (Fig. 1A). Diseased trees with decayed roots may then show symptoms of foliar chlorosis, thinning leaves, defoliation, and eventually decline within a few months to several years. The damaged and fragile roots (Fig. 1B) also make the trees easily toppled over by strong winds and heavy rains. Basidiocarps are occasionally formed on trunks of infected trees (Fig. 1C-D). The sexual reproduction system of *P. noxius* has remained unclarified, partly due to the lack of clamp connections for diagnosing compatibility (P. Ann et al., 2002).

**Fig. 1.**
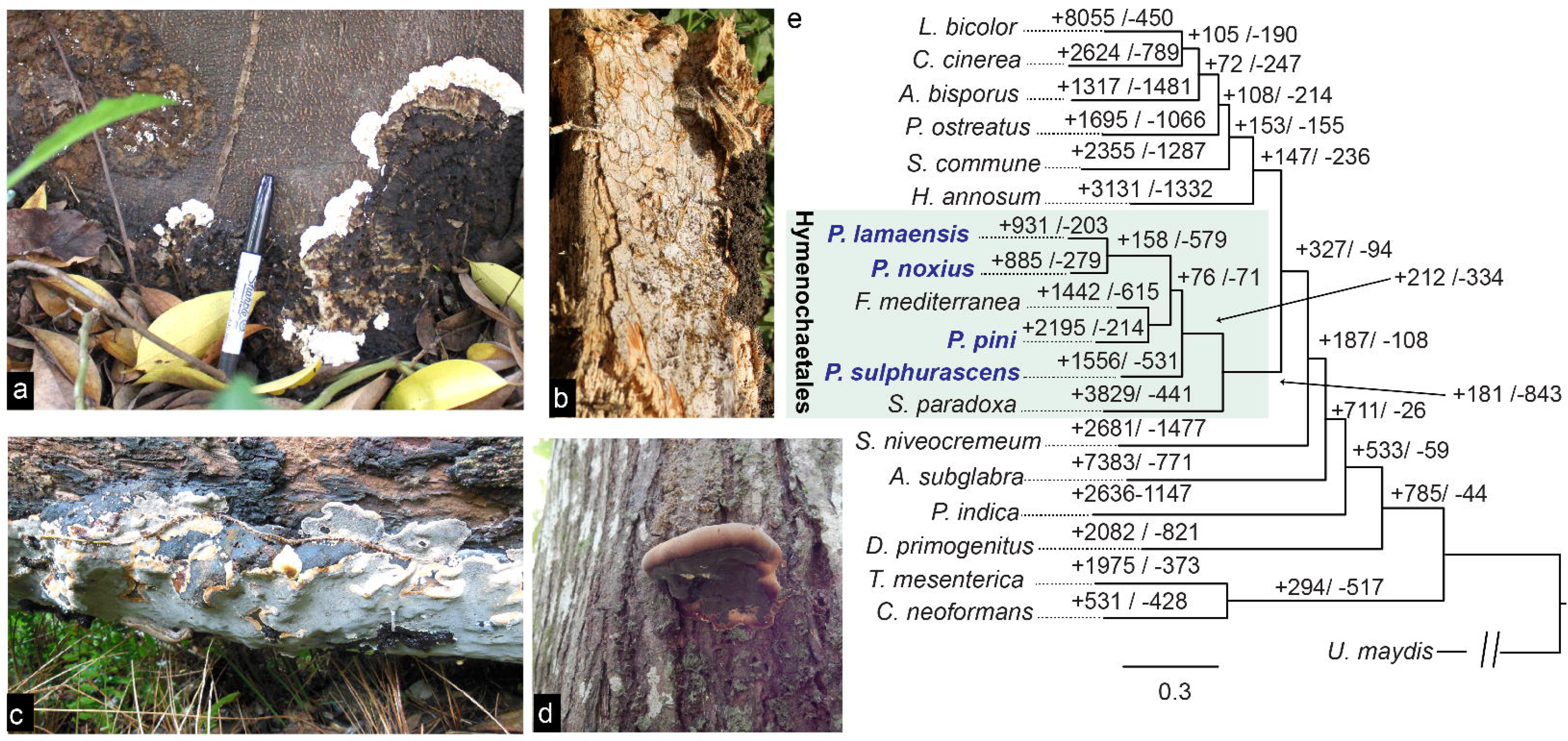
Life stages of *P. noxius* and comparative genomic analysis of Hymenochaetales species. (a) The mycelial mat with young creamy leading front and aged brown section. (b) In advanced stage of decay, the hyphae form a network of brown zone lines permeating the soft and white wood tissue. (Lower left and lower right) (c and d) Basidiocarps are perennial and can be resupinate (c) or grow into a sessile bracket-like conk with a broad basal attachment (d). The distinctive greyish-brown surface is the hymenial layer with irregularly polygonal pores, containing four-spored basidia, ellipsoid and hyaline basidiospores, but no hymenial setae. (e) The phylogeny of four *Phellinus* species with 15 other species of Basidiomycota based on a concatenated alignment of single-copy orthologous genes. All nodes have 100 out of 100 bootstrap replicates. The numbers of gained (“+”) and lost (“-”) gene families along each branch of the phylogeny is annotated.

Brown root rot disease caused by *P. noxius* is widespread in tropical and subtropical areas in Southeast and East Asia, Oceania, Africa, Central America and the Caribbean. The geographical distribution appears to be related to the growth temperature range of *P. noxius*: 10-12°C to 36°C, with optimum growth at 30°C. In the past 20 years, brown root rot disease has become a serious threat to a variety of perennial fruit trees, ornamental and landscape trees, and shade trees in Taiwan (Chung et al., 2015) and in Ryukyu and Ogasawara Islands of Japan (Akiba et al., 2015). In Australia, it occurs in the natural and commercial plantation forests and orchards along the east coast, and has killed many trees in the Greater Metropolitan area of Brisbane Queensland (Schwarze, Jauss, Spencer, Hallam, & Schubert, 2012). In West Africa and China, brown root rot was reported as the most devastating root disease attacking the rubber plantations (Nandris et al., 1987). Trees in urban areas and plantation forests in Hong Kong and Macao have also been seriously affected (Huang, Sun, Bi, Zhong, & Hu, 2016; Wu et al., 2011). Field observations support the root-to-root spread as a major transmission mode of the epidemic. Recent population genetics studies based on SSR markers revealed highly diverse genotypes within populations and nearly identical genotypes from neighboring infected trees, suggesting that *P. noxius* spreads over short distances via root-to-root contact of the hosts, and the genetically variable basidiospores are likely involved in long-distance dispersal and the establishment of unique clones in new disease foci (Akiba et al., 2015; Chung et al., 2015).

In the present study, we aimed to further understand the evolution, reproductive system, and epidemiology in *P. noxius*, on the whole genomic basis. To achieve this, we first sequenced, assembled and annotated the genome sequences of four species from Hymenochaetales: *Phellinus noxius*, *Phellinus lamaensis* (wood decomposing fungus that causes a white pocket rot only on dead heartwood trees), *Phellinus sulphurascens* (syn. *Coniferiporia sulphurascens* (Zhou, Vlasák, & Dai, 2016); pathogen responsible for laminated root rot in Douglas-fir/ true fir), and *Porodaedalea pini* (syn. *Phellinus pini;* trunk pathogen of conifers). Focusing on *P. noxius*, its ~31Mb genome was sequenced to a high level of completion containing telomere-to-telomere chromosome sequences. By comparing against representative species of Basidiomycota, we investigated the genomic conservations and specialisations of this group and how these features possibly relate to the lifestyle of a wood-decayer. Second, we collected *P. noxius* isolates from diseased trees across Taiwan and Japanese offshore islands in 2007-2014. We sequenced the whole-genomes of these 60 isolates and realigned against the *P. noxius* reference genome to identify single nucleotide polymorphisms (SNPs). Based on the genetic variation, the phylogenetic relationship of *P. noxius* populations within this Asia Pacific region was determined. We also quantified the extent of intra-isolate polymorphism to infer the frequencies of morphologically indistinguishable mono- and heterokaryons at infection sites. This allowed the confirmation of heterokaryosis is not necessary for pathogenicity in *P. noxius*.

## Methods

### Strain preparation and sequencing

Genome sequencing of three *Phellinus* species and *P. pini* (Text S1) was performed using both Pacific Biosciences (*P. noxius*, *P. lamaensis, P. sulphurascens* and *P. pini*) and Illumina (*P. noxius*) platforms. DNA was isolated using the CTAB method (Chung et al., 2015). At least 15 μg DNA was used for a 20 kb library prep according to the manufacturer’s instructions. Sequencing was performed on a Pacific Biosciences RS II system using 8 SMRT cell per run, P6C4 chemistry and 360 min movie time. A total of 5-7 SMRT cells were run per species yielding a raw depth of coverage of 173-266X. For samples underwent *de novo* assembly, genomic DNA were sheared in M220 Focused-ultrasonicator™ (Covaris) in microTUBE-50 (Covaris) under 250bp program (duty factor 20%, treatment for 120 sec) with gel size selection after adaptor ligation. For population resequencing samples, genomic DNA were sheared in microTUBE-130 (Covaris) using 500bp shearing program (duty factor 10%, treatment for 60 sec). Genomic libraries were prepared using TruSeq DNA LT Sample Prep Kit (Illumina). Input of 1 μg sheared DNA was used for end-repair, A-tailing, adaptor ligation, and gel size selection. Size range at 600-700bps range was selected from gel and amplified by 5 cycles of PCR. In addition, Illumina mate-pair libraries were generated using 8 μg of genomic DNA with gel size selection of tagmented DNA at 2-4 kb, 4-7 kb, 7-10 kb, and 10-20 kb, and amplified by 10~15 cycles of PCR. These libraries were normalized by KAPA Library Quantification Kit (KAPA Biosystems), and pooled equally for PE2*300 sequencing on MiSeq V2 sequencer. The descriptions of the raw genomic data are available on Table S1.

To aid annotation for each of the species in this study, 7- to 21-day old mycelia from PDA cultures were used for RNA-seq. RNA-seq libraries were constructed using the Illumina TruSeq Stranded mRNA HT Sample Prep Kit with the dual index barcoded adaptors. Input of 3 μg of total RNA was used for each sample for two rounds of oligo-dT bead enrichment, and the ligated cDNA were amplified by 10 cycles of PCR. The Stranded mRNA libraries were quantified by Qubit and molar concentrations normalized against the KAPA Library Quantification Kit (KAPA Biosystems) for Illumina platform. The transcriptome libraries were pooled at equal molar concentrations, and PE2*151nt multiplexed sequencing was conducted on HiSeq 2500 sequencer. The descriptions of the raw RNA-seq data are available on Table S2.

### Nuclear quantification

Nuclei in the growing hyphal tips were stained following the procedure described by Chung et al. (2015). The mycelium on the slide was mounted in 20 μl of a DAPI (4',6-diamidino-2-phenylindole) solution (10 μg/ml in ddH_2_O) for an hour, destained in ddH_2_O for 30 min, then observed under a OLYMPUS BX41 microscope (Shinjuku-ku, Tokyo, Japan) equipped with filter cube U-MWU2 (BP 330–385 nm, LP 420 nm). Images were captured by using a Canon (Ohta-ku, Tokyo, Japan) digital camera EOS 700D. One hundred hyphal cells per strain were counted.

### Genome assembly

Genome assembly of different species was carried out with Falcon (ver 0.5.0; Chin et al., 2016) and were improved using Quiver (Chin et al., 2013) and finisherSC (Lam, LaButti, Khalak, & Tse, 2015). For assembly of individual strains of *P. noxius*, Illumina paired end reads were trimmed with Trimmomatic (ver 0.32; options LEADING:30 TRAILING:30 SLIDINGWINDOW:4:30 MINLEN:50; Bolger, Lohse, & Usadel, 2014) and subsequently assembled using SPAdes (ver 3.7.1; Bankevich et al., 2012). Multiple mate-pair reads were available for three strains of *P. noxius* (KPN91, A42 and 718-S1) and they were assembled using ALLPATH-LG (ver 49688; Butler et al., 2008) assembler and improved using Pilon (Walker et al., 2014). The *P. noxius* assembly was further merged with metassembler (ver 1.5; Wences & Schatz, 2015), misassemblies were identified using REAPR (ver 1.0.18; Hunt et al., 2013) and manually corrected.

### Gene predictions and functional annotation

For *P. noxius*, the gene predictor Augustus (ver3.2.1; Stanke, Tzvetkova, & Morgenstern, 2006) was trained on a gene training set of complete core genes from CEGMA (ver2.5; Parra, Bradnam, & Korf, 2007) and subsequently used for manual curation of ~1000 genes. Annotation was then run by providing introns as evidence from RNA-seq data. For *P. lamaensis*, *P. sulphurascens* and *P. pini*, genes were predicted using Braker1 (ver 1.9; Hoff, Lange, Lomsadze, Borodovsky, & Stanke, 2016) pipeline that automatically use RNA-seq mappings as evidence hints and retraining of GeneMark-ES (Borodovsky & Lomsadze, 2011) and Augustus. Gene product description was assigned using blast2go (ver 4.0.7; Conesa et al., 2005) and GO term assignment were provided by ARGOT2.5 (Lavezzo, Falda, Fontana, Bianco, & Toppo, 2016). The web server dbCAN (HMMs 5.0, last accessed September 5 2016; Yin et al., 2012) was used to predict CAZymes from the protein sequences of all species, while AntiSMASH (ver 3.0; Weber et al., 2015) was used to predict secondary metabolite gene clusters. For dbCAN results, only hits with <= 1 x 10e-5 e-value and >= 30% HMM coverage were considered, while overlapping domains were resolved by choosing hits with the smallest *P*-value. Proteome completeness were assessed with BUSCO (ver 2.0; Simão, Waterhouse, Ioannidis, Kriventseva, & Zdobnov, 2015) using the Basidiomycota dataset.

### Comparative genomics analysis

Gene families were determined by OrthoFinder (ver 1.0.6; Emms & Kelly, 2015). Then, MAFFT (ver 7.271; Katoh & Standley, 2014) was used to align sequences in each of 1,127 single-copy orthogroups. Alignments results with less than 10% alignment gaps were concatenated, and the outcome was taken to compute a maximum likelihood phylogeny using RAxML (ver 7.7.8; Stamatakis, 2006) with 100 bootstrap replicates. Gene family gain and loss in different positions along the global phylogeny leading to *P. noxius* were inferred using dollop (Felsenstein, 2005). Pfam and GO enrichments were carried out on these gene families using TopGO (ver 2.10.0; Alexa & Rahnenfuhrer, 2016). Sequences to be included in the phylogenetic tree for NACHT domain containing genes were selected on the basis of the presence of pfam domain PF05729.

### MAT locus

Homologs of the conserved genes in mating locus *A* and *B* were annotated in *P. noxius*, *P. pini*, *P. sulphurascens* and *P. lamaensis* (e-value < 10^−5^) then subjected to InterProScan 5 (ver 5.20-59.0; Jones et al., 2014) and Pfam (ver 30.0; Finn et al., 2016) for protein signature prediction. Syntenic alignment of *A* locus among *Phellinus* spp. and other species (sequences/annotations retrieved from Joint Genome Institute MycoCosm (Grigoriev et al., 2014)) was plotted using genoPlotR package (Guy, Roat Kultima, & Andersson, 2010) in R. The sequences of *A* locus in KPN91, 718-S1 and A42 were further compared by MUMmer (ver 3.20; Stefan Kurtz et al., 2004) and PipMaker (http://pipmaker.bx.psu.edu/cgi-bin/pipmaker?basic; Schwartz et al., 2000). For identification of candidate pheromone genes, all the potential open reading frames were filtered for small peptides with C-terminal CaaX motif, then searched for pheromone homologies against Pfam-A and scanned for farnesylation signal by PrePs (http://mendel.imp.ac.at/PrePS/; Maurer-Stroh & Eisenhaber, 2005).

The sequences of *HD* and *STE3* genes were analyzed for 10 single-basidiospore isolates originating from a basidiocarp by the dideoxy termination method (primers listed in Table S3). Long-range PCR followed by primer walking was performed to sequence the highly diverse regions containing *HD1*-*HD2* in *A* locus. The outer primers were designed manually based on the alignment of all the isolates; the inner primers were developed step-by-step according to previous sequencing results. The PCR reaction was performed in 30-μl reaction mixture containing 50 to 100 ng genomic DNA, 0.2 mM dNTP, 1X Ex Taq buffer [proprietary, containing 20mM Mg^2+^] (Takara Bio Inc., Japan), 0.67 μM forward and reverse primers, and TaKaRa Ex Taq® DNA polymerase (Takara Bio Inc., Japan). The thermal cycling parameters were 1 cycle of 95°C for 3 min, 30 cycles of 95°C for 30 s, 54°C for 30 s, and 72°C for 60–270 s (according to different product sizes, ~2 kb/min), and a final extension step of 72°C for 10 min. The PCR products were sequenced by Genomics Biotechnology Inc. (Taipei, Taiwan). DNA trace data were visualized using 4Peaks (http://nucleobytes.com/4peaks) and assembled using DNA Sequence Assembler in DNA Baser (http://www. dnabaser.com).

### Population genomics

Paired end reads of 60 *P. noxius* strains (description listed in Table S4) were aligned to the KPN91 reference using Smalt (ver 5.7; www.sanger.ac.uk/resources/software/smalt/). Removal of PCR duplicates and bam file sorting were implemented with Picard (http://broadinstitute.github.io/picard/) and samtools (ver 1.3-20-gd49c73b; Li et al., 2009). The first round of variant identification was implemented in Varscan (ver 2.4.0; Koboldt et al., 2012) and degree of heterokaryon was inferred in each strain based on allele frequency and total number of heterozygous SNPs called. The final list of SNPs was ascertained by combining evidences from samtools (ver 1.3-20-gd49c73b; Li et al., 2009) and FreeBayes (ver 1.0.2-16-gd466dde; Garrison & Marth, 2012). A maximum likelihood phylogeny of the SNPs segregating in these 60 strains was produced using FastTree (Price, Dehal, & Arkin, 2009). Plink (ver 1.9; C. C. Chang et al., 2015) was used to subset biallelic SNPs without linkage (filtering options: – maf 0.05 – indep-pairwise 50 5 0.2), which were clustered using fastSTRUCTURE (ver 1.0; Raj, Stephens, & Pritchard, 2014) to determine the optimal number of populations. Strains were phased using samtools (ver 1.3-20-gd49c73b; Li et al., 2009) and one haplotype was chosen at random. Consensus sequences were generated from each strain using bcftools (ver 1.3.1; Danecek et al., 2011) and population genetics analyses were conducted using Variscan (ver 2.0.3; Vilella, Blanco-Garcia, Hutter, & Rozas, 2005).

## Results

### Genomes and annotations of four Hymenochaetales members

We produced a 31.6 Mb *Phellinus noxius* genome reference assembly from a Japanese KPN91 isolate combining both Pacific biosciences and Illumina sequencing platforms (Methods). The nuclear genome of *P. noxius* is assembled into 12 scaffolds with six assembled into chromosomes from telomere to telomere, while the mitochondrial genome is assembled into a single sequence of 163.4 kb. For a comprehensive understanding of genome evolution amongst members of the hymenochaetoid clade, we also sequenced and assembled three additional species: *P. sulphurascens, P. lamaensis* and *P. pini*, as well as two more isolates (A42 and 718-S1) of *P. noxius*. The three assemblies of *P. noxius* have N50s of 2.4-2.7 Mb, whilst the other three genome assemblies comprise 30.7-53.3 Mb with N50’s of 570 kb-2.7 Mb.

A total of 9,833-18,103 genes were predicted in each species, which are 82-94% complete (Table S5). To compare these predicted proteins to those of other basidiomycetes to explore chromosome and gene family evolutionary dynamics, we selected the proteomes of fifteen species that are highly finished from the 1000 Fungal Genomes Project (Table S5). The Hymenochaetales species have median intergenic and intron lengths of 507-634 bp and 59-60 bp, respectively, which are comparable with those observed in genomes of other basidiomycetes. The maximum likelihood phylogeny based on 1,127 single-copy orthologs placed these species with *Fomitiporia mediterranea* and *Schizopora paradoxa*, two other species of the Hymenochaetaceae group with strong bootstrap support (Fig. 1E). This phylogenetic relationship is consistent with previous findings (Larsson et al., 2006), and species with similar genome sizes and pathogenic habits are grouped together. For instance, *P. noxius* and *P. lamaensis* with compact genome sizes of ~31 Mb are grouped together, while the trunk rot pathogens *P. pini* and *F. mediterranea* show an expansion of their genome sizes to 53-63 Mb and are also grouped together.

To explore the genomic architecture underlying the biological attributes of Hymenochaetales, we sought to identify genes and protein domains specific to Hymenochaetales by determining when a new gene family arose and if the family has expanded or contracted. In total, 7,125-11,659 proteins in the Hymenochaetales order are clustered together in 5,184 families. Acquisition of gene families was mainly found at the tips of the phylogeny (531-8,055 families) suggesting each species has a repertoire of specific genes. The seven Hymenochaetaceae species have a total of 62 enriched protein domains compared to other basidiomycetes (Fig. S1). Gain of domains are highlighted such as fungal specific transcription factors (Fungal_trans; 53.8 *vs* 41.6 copies) and peroxidase associated protein domains (Peroxidase_ext; 16.7 *vs* 3.9 copies). These are expected as they are required for efficient degradation of lignin, a tough biopolymer present in woody plants (Dashtban, Schraft, Syed, & Qin, 2010). Consistent with this, CAZymes spanning eight families of lignin-degrading enzymes (AA1-AA8) which include laccases, peroxidases, oxidases, and reductases, and a lytic polysaccharide mono-oxygenase family (LPMO; AA9) were found in all Hymenochaetales species (Table S6).

### Genome conservation and specialisation of Phellinus noxius

Although *P. noxius* has a compact genome, 74% (7,313) of its 9,833 predicted genes have orthologs from at least one of the other basidiomycetes, suggesting most of the basidiomycetous core genes are conserved. As a chromosome level assembly is now available in Hymenochaetales, we attempted to characterize chromosome architecture and evolution amongst the Agaricomycotina sub-division. The number of known karyotypes in Basidiomycota ranges from 11 to 14 chromosome pairs (Table S5), which suggests a possible ancestor with similar chromosome numbers. We constructed a linkage network of seven selected species with chromosome sequences based on single-copy orthologs between species pairs. Indeed, we identified 13 distinct linkage groups (LG) providing strong evidence that chromosome macro-synteny is largely conserved since the common ancestor of Agaricomycetes (Fig. 2A). Strikingly, such a relationship even extended to Dacrymycetes where multiple scaffolds can be predominantly assigned to different linkage groups, but it is no longer apparent when compared to Tremellomycetes. Certain scaffolds are found to connect different linkage groups, implying inter-chromosomal rearrangements. For example, *P. noxius* scaffold1 is strongly clustered in LG11 but also shows linkage to LG10, implying a translocation from scaffold5 (Fig. 2). Within each linkage group, gene collinearity is no longer apparent, suggesting high levels of intra-chromosomal rearrangements which have been observed in different fungal groups (Hane et al., 2011) (Fig. 2B).

**Fig. 2.**
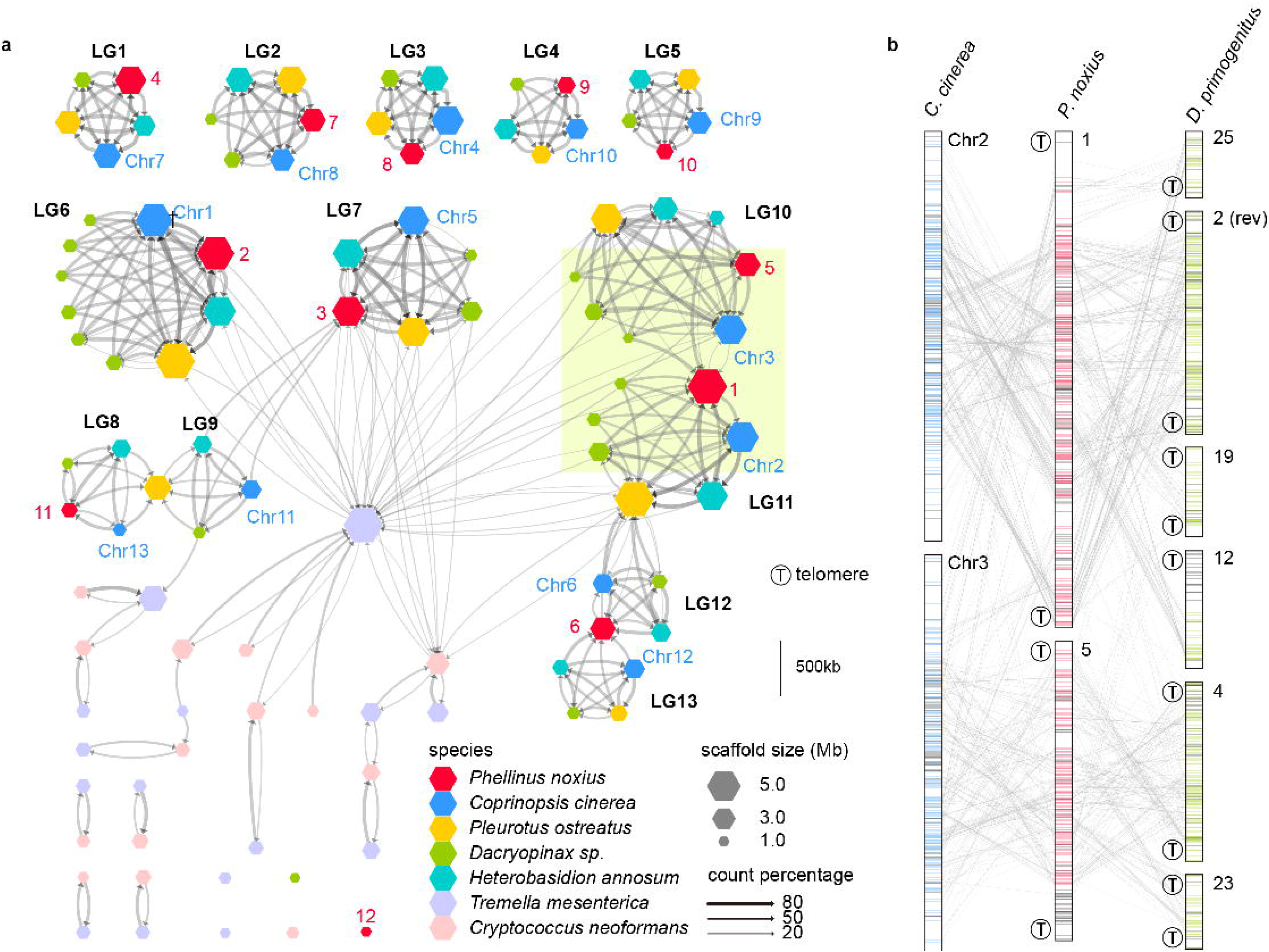
Linkage group (LG) network of Basidiomycota. (a) The linkage groups were identified by linking single-copy orthologs of scaffolds between species pairs. All scaffolds included in this plot are larger than 500 kb. Each directional edge points toward the reference chromosome. Edges were weighted by the number of single-copy orthologs, and an edge was filtered if it has a weight smaller than 20 or less than 40% of the sum of all weights out of a node. (b) Cross-mapping of single-copy orthologs in LG10 and 11. Scaffold names are shown at the upper-right side of each sequence, with detectable telomeric regions labelled as ‘T’ at the upper-left side.

Consistent with the fact that *P. noxius* is an extremely fast grower (P. Ann et al., 2002), we identified a strikingly 7-fold increase in 1,3-beta-glucan synthase (14 compared to an average of 1.8 copies), which is responsible for the formation of beta-glucan components in fungal cell wall (Douglas, 2001). *P. noxius* contains a comprehensive repertoire of carbohydrate-active enzymes; a total of 416 proteins of its proteome were identified as CAZymes (Table S6). This number makes up 4.23% of *P. noxius*’s proteome which is more than those in the other Hymenochaetales, suggesting these genes are necessary and have been retained despite a genome size compaction. Taken together, being able to grow fast and the diversity of the CAZymes encoded in the *P. noxius* genome suggest its capability to infect a wide range of hosts. Additionally, counts of WD40 protein domains are highest in *P. noxius* and *P. lamaensis* despite their small genome size (Table S7). This domain is important in protein-protein interactions of cellular networks and is usually associated with additional domains (Leipe, Koonin, & Aravind, 2004). Interrogating this expansion revealed the association of WD40 domains with the AAA and NACHT domains (Table S7), both of which are NTPase domains and such combinations are commonly found in nucleotide-binding-oligomerization-domain like receptors (NLRs). A maximum likelihood phylogeny of the NACHT domain proteins shows different expansions of NLR subfamilies in fungi (Fig. S2). In particular, the C2-NACHT-WD40(n) subfamily has only been found exclusively in a few basidiomycetes (Van der Nest et al., 2014) and is present in the highest copy number in *P. noxius* and *P. lamaensis*. Other expansions include UDP-glucuronosyltransferases, which catalyse conjugation reactions in the biotransformation of xenobiotics released by its host environment (Jancova, Anzenbacher, & Anzenbacherova, 2010).

### P. noxius displays a bipolar heterothallic mating system

Determination of mating loci and reproductive mode is considered high-priority in fungal genome analyses as they are the primary determinants of how a fungus expands and generates diversity. Mate recognition of sexual reproduction in Basidiomycota is known to be controlled by two unlinked loci, named as *A* and *B* locus. A conserved head-to-tail orientation of *HD1-HD2* in *HD* pair 1 was found in Hymenochaetales (Fig. 3) which is different from in most species in Agaricomycetes (James et al., 2013). Alignment of 100-kb sequences upstream and downstream of *A* locus in *P. noxius* isolates KPN91, 718-S1 and A42 revealed that *A* locus is highly polymorphic (*HD* pairs in particular) despite well-conserved flanking regions (Fig. S3). For *B* locus, only one *STE3* encoding seven transmembrane helices was identified; four pheromone genes were identified but not physically linked to the highly monomorphic *STE3* (Fig. S4, Table S8, Text S1), which is characteristic of a bipolar mating system.

**Fig. 3.**
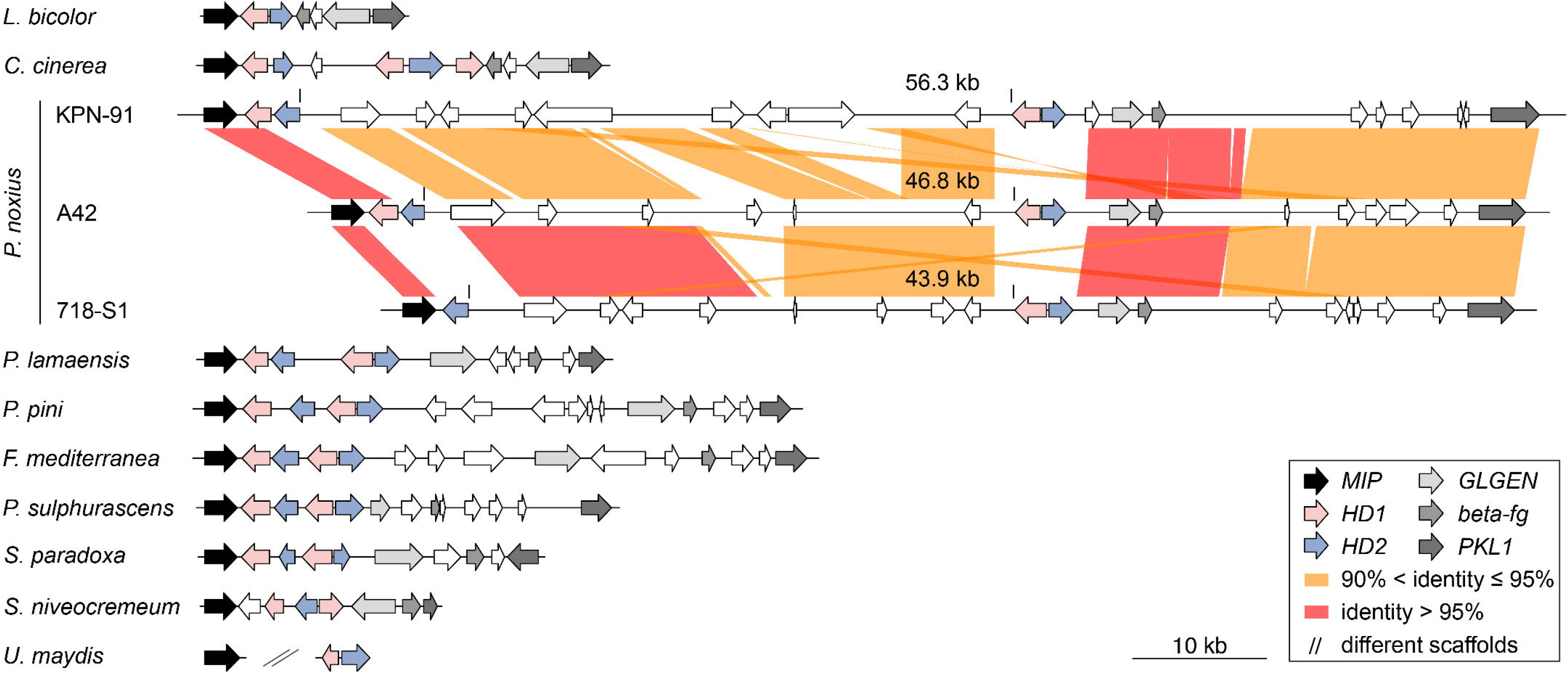
Synteny around the *A* mating locus. The analysis included *Phellinus noxius, Porodaedalea pini, Phellinus sulphurascens, Phellinus lamaensis*, two other species from Hymenochaetales species (*Fomitiporia mediterranea, Schizopora paradoxa*), two species from Agaricales (*Laccaria bicolar, Coprinopsis cinerea*), *Sistotremastrum niveocremeum*, and *Ustilago maydis*.

To further confirm this observation, allele diversity was analyzed by resequencing the *A* and *B* loci from 10 single-basidiospore isolates originating from a single basidiocarp (Table S9). Sequencing of *STE3* revealed two highly similar alleles (b1, b2), with 99.5% amino acid identity. The only variant (244V/A) is considered a semi-conservative mutation (valine and alanine are nonpolar aliphatic acids) and may have minor or no effect on protein function. Previous studies have shown that although STE3 is not involved in mating type determination in bipolar fungi, variations can still be observed (James, Lee, & van Diepen, 2011; James, Srivilai, Kues, & Vilgalys, 2006; Niculita-Hirzel et al., 2008). Primer walking of *A* locus revealed two distinct haplotypes which suggests a heterothallic bipolar reproductive mode in *P. noxius* (Fig. S5): the a1 and a2 alleles of *HD* pair I shared an overall 78% nucleotide identity; the two alleles of *HD* pair II were highly divergent and the HD1 domain in a2 allele contains a 1-bp and a 9-bp deletions and has become a pseudogene. The loss of HD1 domain was also found in the *HD* pair I of 718-S1 (Fig. 3), suggesting that at least one of the HD1 motifs in *A* locus is dispensable for a functional HD1-HD2 heterodimer in *P. noxius*. The presence or absence of specific HD domains reflects phylogenetic characteristics and has been commonly observed in fungi (Kües, 2015).

### P. noxius contains a highly polymorphic A locus

Distinct from all the other fungi, *P. noxius* has an exceptionally highly expanded *A* locus across a ~60-kb region (Fig. 3). There are two pairs of HD1-HD2 gene located ~50-kb away from each other. In the case of *P. noxius* KPN91, 14 genes were annotated at the *A locus* (Fig. 3). Orthology analysis placed these genes into 9 families that are also present in other fungi (Fig. 4). However, in the genomes of other species, the majority of homologous members of different families are not located in proximity to each other (Fig. 4). This indicates that the large interval in *P. noxius* is a result of accumulating transposition of genes from conserved fungi families not in proximity to *A* locus (Fig. 4). Interestingly, different *P. noxius* isolates display different interval lengths at *A* locus, suggesting that transposition events may be frequent (Fig. 3). Whether transposition plays an additional role in the mating system remains to be clarified.

**Fig. 4.**
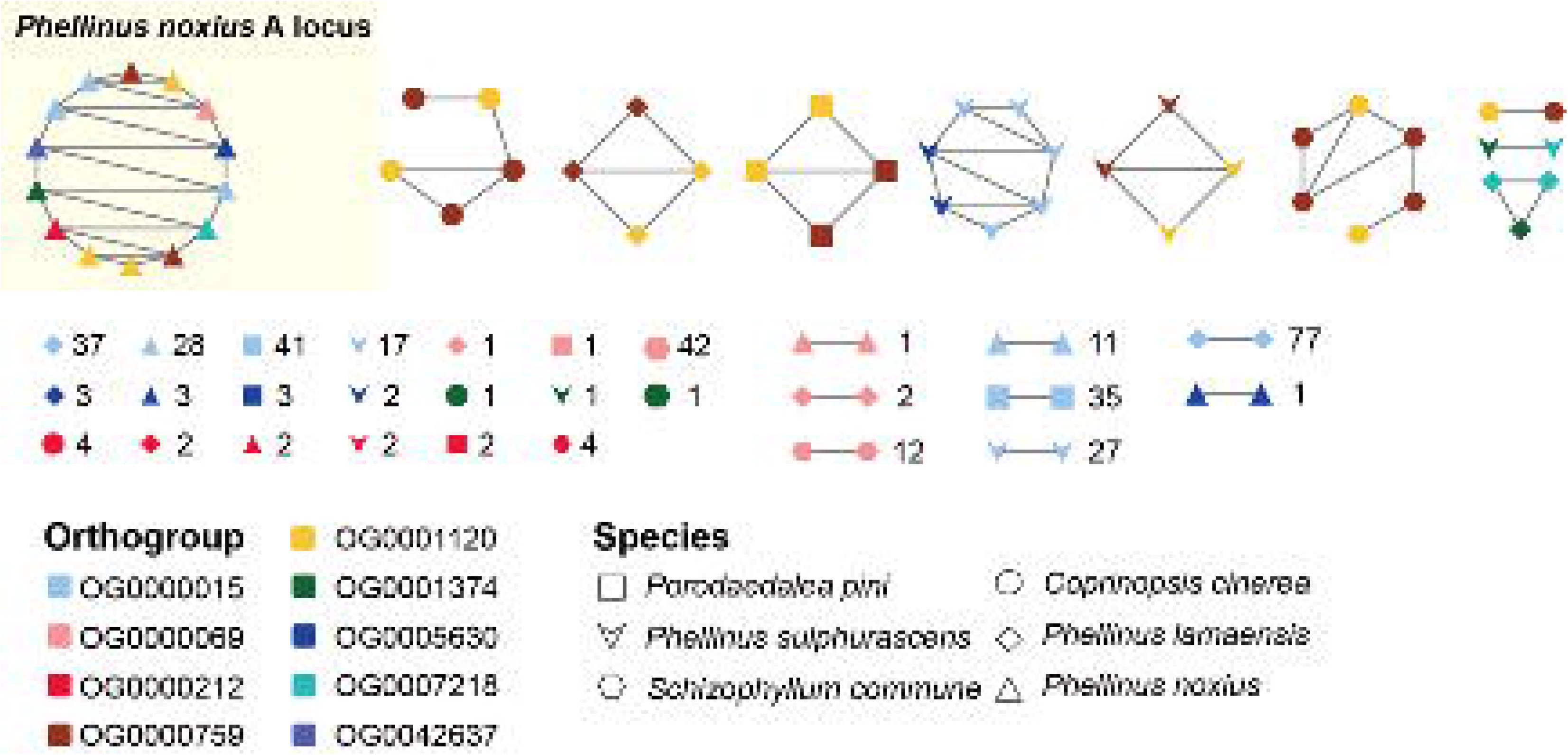
Orthology network of genes in the *A* locus of *P. noxius*. Each node in this plot represents a gene. An edge is added if two genes are in proximity (physically separated by less than two genes). Numbers next to the edges and nodes are number of occurrences of different combinations of genes in each species. For example, 77 members of OG0000015 orthogroup in *P. lamaensis* are found adjacent to each other on the genome. Another 37 members of this orthogroup are dispersed throughout the genome and are not located in proximity to any members of the 9 orthogroups.

### Sequence analysis of 60 P. noxius strains

To further understand the regional dissemination of *P. noxius*, we sequenced the genomes of 60 isolates (Fig 5A) originating from diseased trees across 13 Taiwan and Japanese offshore islands from 2007-2014. This collection was sequenced to a median depth of coverage of 35X. To characterize chromosomal synteny relationship, additional mate pair libraries were sequenced for two isolates (718-S1 and A42). An average of 96% mapping rate was achieved after aligning reads from each strain to the KPN91 reference genome. The descriptions and statistics of the strains are provided in Table S4.

**Fig. 5.**
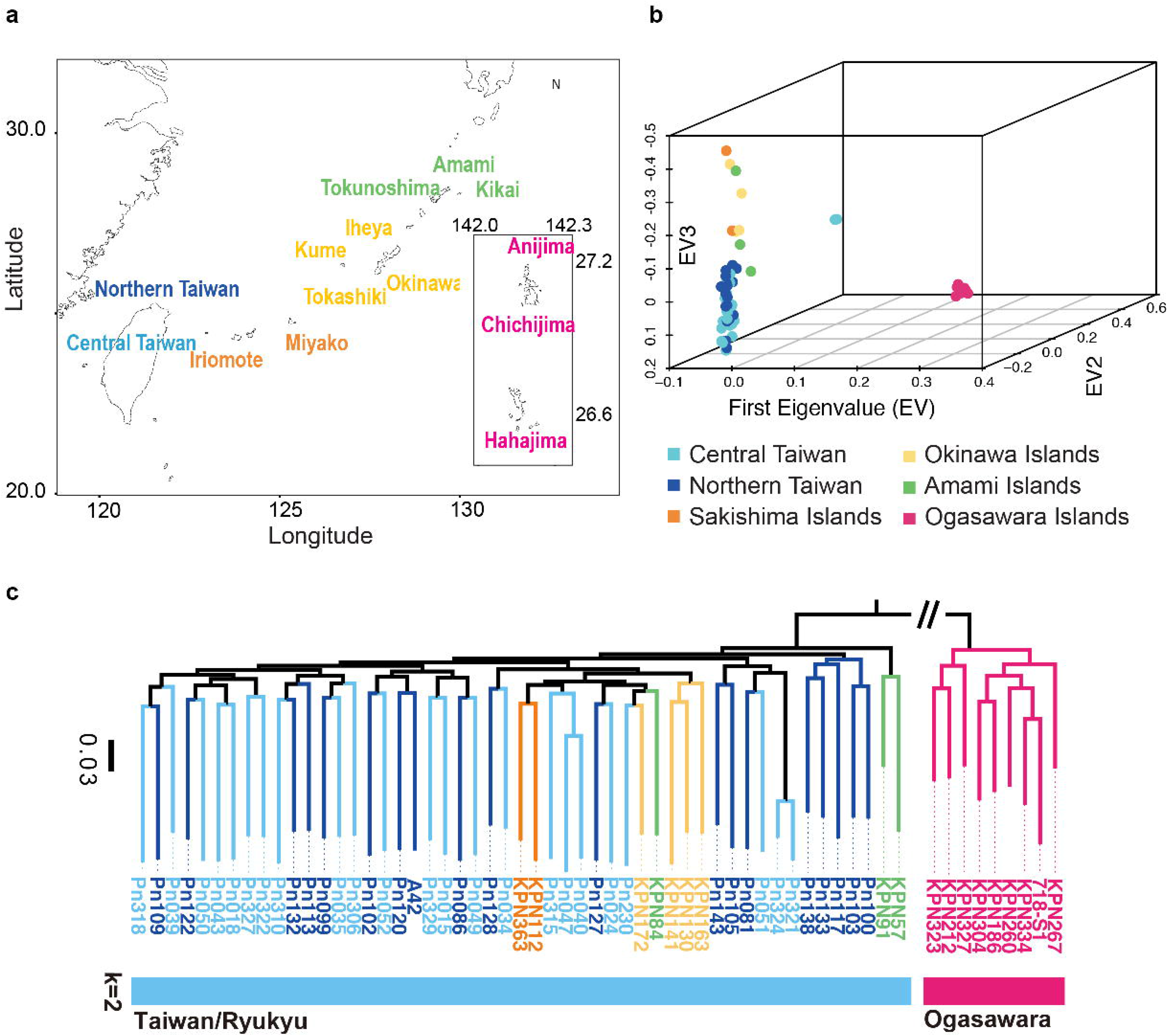
The population genomics of 60 *P. noxius* isolates. (a) Map of Taiwan and offshore islands of Japan showing origins of the 60 sampled *P. noxius* isolates. Ogasawara islands were conveniently drawn below Ryukyu islands so this does not represent their actual location. (b) PCA plot of 60 *P. noxius* isolates using genome-wide variation data sampled from 13 islands by the first three eigenvectors. (c) Top: Phylogenetic tree with 100 bootstrap using SNPs computed from alignment to KPN91 reference. Point separating the Taiwan-Ryukyu and Ogasawara island isolates was used as root. Nodes with >90% bootstrap were labelled with circles. Bottom: fastSTRUCTURE analysis of the linkage independent pruned set of variation data. A model with two ancestral components (*K*=2) had the highest likelihood to explain the variation of genome-wide structure on the 60 isolates. Also see Fig. S7 for different *K*.

### Diversification of P. noxius across Pacific Ocean islands

We hypothesized that there may be three populations segregating in this area of Pacific Islands: Taiwan, Ryukyu and Ogasawara islands. To examine the population structure of *P. noxius* in this area, we performed principal-component analysis (Fig. 5B) on the SNP variants from individual phased haploid genomes. The major principal component divided Ogasawara islands samples from the rest of the samples, which are located geographically 1,210 km apart. The second and third principal components defined a tight cluster of Taiwanese isolates with considerable overlaps between cities despite being 170 km apart. Most isolates from Ryukyu islands can be differentiated from those from Taiwan (Fig. S6). Interestingly, we identified three isolates from Taiwan that are distinctive from both the Taiwan-Ryukyu and Ogasawara island clusters, suggesting a possibility of more genetically distinct populations within the Taiwan island.

We constructed a maximum likelihood phylogeny based on 1,837,281 high confidence SNPs of *P. noxius* (Fig. 5C). Concordant with the PCA, the nine isolates from Ogasawara islands form a distinctive lineage, while the 51 Taiwanese and Ryukyu islands isolates are grouped together forming another major lineage. We inferred the population structure in 144,426 unlinked bialleic sites using the Bayesian model-based clustering approach implemented by fastSTRUCTURE (Raj et al., 2014). Consistent with the phylogeny, two Taiwan-Ryukyu and Ogasawara lineages were identified and circulating in this region (highest likelihood with *K =* 2; Fig. 5C). Higher *K* values shows that the isolates from Ogasawara islands remain one cluster, while the rest of the isolates (independently of their geographical origins) are grouped into genetically homogeneous clusters with little admixture (Fig. S7).

Further inspection of the phylogeny indicated the possibility of gene flow in this region despite physical separation by the sea. Within the Ogasawara clade, the isolates can be further grouped by their geographical origin (two main islands: Hahajima and Chichijima island separated by 50km) with the exception of KPN334 strain. Strain KPN260 was collected from Anijima island which is geographically in proximity to Chichijima island but was grouped with the Hahajima island isolates. Strains collected from within Amami and Okinawa islands are not grouped together on the phylogeny, suggesting independent origins of *P. noxius* infection in both regions. The pattern of gene flow was more apparent in the 42 isolates collected in two cities of Taiwan, where the isolates were not grouped in the phylogenetic tree according to their geographical origins.

### Stable structural variation and intra-isolate polymorphism

The detection of frequency of heterokaryons and polyploids of *P. noxius* in nature remain challenging as its arthospores contain 1-5 nuclei (based on quantification of 145 arthospores from A42 in this study) and multiple allelic fragments in SSRs (2-4 alleles per locus) are commonly found in populations (Akiba et al., 2015; Chung et al., 2015). All the strains were isolated from either a single arthospore or fungal mat, and thus allowed for the analysis of variation in genome structure. We found no deviation in coverage across every scaffold in all strains (Fig. S8), suggesting a stable number of chromosome copies in *P. noxius*. To distinguish the frequency of mono- or heterokaryons in nature, heterozygous SNPs and minor allele frequency (MAF) distribution were inferred in each strain (Fig. 6). We identified four groups (A to D) that clearly differ in heterozygosity and MAF. The group A with the lowest heterozygosity of averaging 0.2 % includes strains 718-S1 and A42, each isolated from a single basidiospore implying they are monokaryotic in nature. All strains in this group exhibited a flat MAF distribution, suggesting that there are spontaneous mutations segregating during the growth of monokaryons that originated at different times. The occurrences of monokaryons are 11% and 47% in the Ogasawara and Taiwan/Ryukyu lineages, respectively. The prevalence of monokaryotic isolates suggests the involvement of basidiospores in disease spread and that heterokaryosis is not required for pathogenicity.

**Fig. 6.**
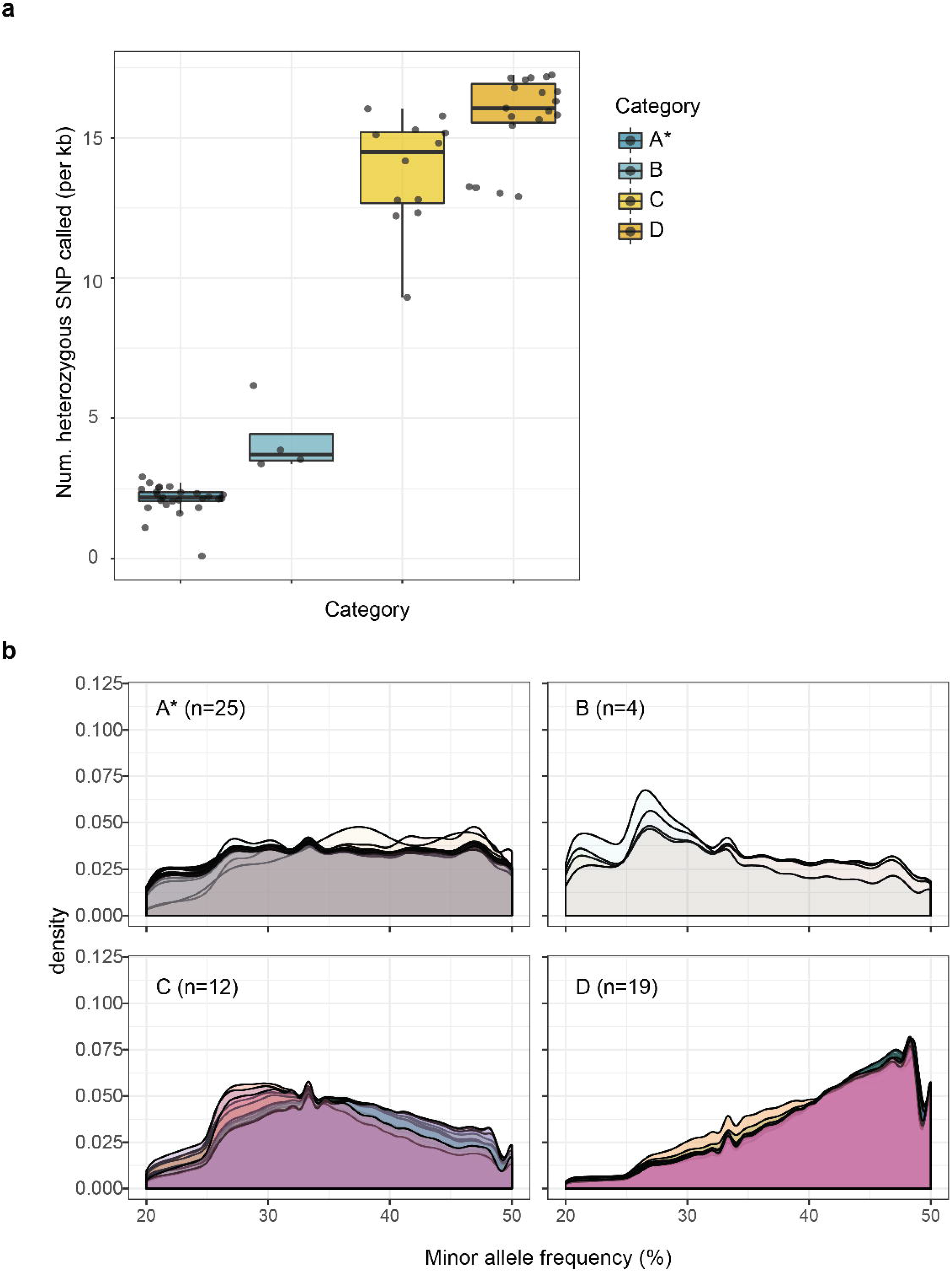
Heterozygosity and allele frequencies of *P. noxius*. (a) Boxplot showing abundances of heterozygous SNPs inferred in each *P. noxius* isolate can be categorised into four groups. (b) Density plot of minor allele frequencies (MAF) of heterozygous SNPs in 60 isolates of *P. noxius*. *Asterisk denote group containing two isolates that were cultured from basidiospores.

58% of strains display a much higher heterozygosity, with strain Pn103 having a 1.7% heterozygosity displaying a peak around 50% in MAF distribution. These strains can be further grouped into three categories with distinct MAF profile and heterozygosities (Fig. 6). The largest group of the three (n=19; group D of Fig.6) has on average 1.6% heterozygosity displaying a peak around 50% in MAF distribution, suggesting the presence of two genetically distinct nuclei within the population, i.e., dikaryons. The remaining two groups did not follow a typical di-karyon MAF distribution, as they peaked around 27%-33% and exhibited different heterozygosities. The deviations may be associated with the number and composition of multiple nuclei in a cell. However, nuclear quantification of seven randomly selected isolates showed no differences between the groups (Fig. S9). These groups may refer to different compositions of two or more genetically distinct nuclei.

### High nucleotide diversity in P. noxius populations

The *de novo* assemblies of three strains of *P. noxius* have on average 97% nucleotide identity and are largely co-linear to each other with apparent genome translocations between Taiwan/Ryukyu (A42 and KPN91) and the Ogasawara (718-S1) isolate (Fig. S10). We quantified sequence diversity using θs and θπ and categorised them into whole genome average, four-fold synonymous sites and replacement sites (Table 1). A large number of segregating SNPs are found in the genome of *P. noxius*, averaging one SNP identified in every 20-59 bp. Nucleotide diversity at synonymous sites θπ-syn is 15.8 and 19.2 per kilobase in Ogasawara island and Taiwan-Ryukyu populations, respectively. This is ~5 fold higher when compared to the majority of species (Leffler et al., 2012) and is likely an underestimate of true diversity as indels were not considered. Taiwan-Ryukyu lineage has a much higher diversity than Ogasawara which is not due to sample size difference; the same was observed when we reanalyzed with only 9 randomly chosen strains in the Taiwan-Ryukyu populations. Extremely high diversity has been reported in natural populations of *Schizophyllum commune* (Baranova et al., 2015). The Tajima’s D is strongly negative in the Taiwan and Ryukyu islands lineage, suggesting an excess of low frequency alleles present in the population possibly as a result of high mutation rate (Baranova et al., 2015) or population expansion. In the Ogasawara population, nucleotide diversity is reduced compared to the Taiwan/Ryukyu lineage and Tajima’s D is positive implying a reduced effective population size; *P. noxius* may have been introduced recently in these islands. Together our data suggest that the two *P. noxius* lineages may have derived from genetically distinct gene pools and have undergone divergent evolutionary scenarios, possibly as a result of different time of introduction, different environments, and human interference in Taiwan-Ryukyu *vs* Ogasawara areas.

**Table 1.** Polymorphism in the two regional lineages of *P. noxius*.

| Population | Analyzed Sites | Segregating sites (S) | Singletons | $\theta\pi$ (x1,000) | $\theta s$ (x1,000) | Tajima's D |
| --- | --- | --- | --- | --- | --- | --- |
| **Whole genome** |  |  |  |  |  |  |
| Ogasawara (n=9) | 30,028,158 | 506,231 | 174,924 | 6.6 | 6.3 | 0.3 |
| TaiwanRyukyu (n=51) | 31,376,691 | 1,538,598 | 545,324 | 8.0 | 10.9 | -0.9 |
| TaiwanRyukyu subset (n=9)<sup>1</sup> | 30,640,276 | 749,681 | 408,021 | 8.1 | 8.9 | -0.5 |
| **Synonymous Sites** |  |  |  |  |  |  |
| Ogasawara (n=9) | 4,651,751 | 182,879 | 59,464 | 15.8 | 14.7 | 0.4 |
| TaiwanRyukyu (n=51) | 4,758,211 | 513,717 | 161,353 | 19.2 | 23.6 | -0.7 |
| TaiwanRyukyu subset (n=9)<sup>1</sup> | 4,704,161 | 269,060 | 136,780 | 19.3 | 20.5 | -0.4 |
| **Replacement Sites** |  |  |  |  |  |  |
| Ogasawara (n=9) | 9,794,069 | 71,497 | 25,981 | 2.9 | 2.8 | 0.2 |
| TaiwanRyukyu (n=51) | 10,018,114 | 230,689 | 93,385 | 3.6 | 5.3 | -1.2 |
| TaiwanRyukyu subset (n=9)<sup>1</sup> | 9,904,211 | 103,907 | 60,313 | 3.6 | 4.0 | -0.6 |

<sup>1</sup> A random of nine isolates were chosen to repeat the analysis.

### Candidates for population differentiation

The Taiwan/Ryukyu strains were mainly isolated from diseased trees in urban settings, while the Ogasawara strains were from trails in natural parks (Table S4). Genomic pairwise F_ST_ revealed moderate differentiation (0.12) between the two genetic lineages of *P. noxius*, which is in accordance with both the phylogeny and fastSTRUCTURE analyses. We identified genomic regions that may contain potential candidates for such environmental origins by investigating 99.9% tail for observed F_ST_ 5-kb windows (Fig. 7). This definition revealed 13 outlier regions. Gene ontology analyses of 42 genes within these regions did not reveal any GO enrichment terms, indicating the sites showing evidence of differentiation may be involved in different functions. Interestingly, the largest region spanning 12 kb at scaffold 6 contains four genes. Homologs from three of the genes are implicated in fungal cell wall organization and fruiting body formation (PNOK_0653900; (Szeto, Leung, & Kwan, 2007)), salt tolerance (PNOK_0654000; (Steffens, Brautigam, Jakoby, & Hulskamp, 2015)), and plant cell wall degradation (PNOK_0654100; xylanase). These are all possible drivers for population differentiation (Apse, Aharon, Snedden, & Blumwald, 1999).

**Fig. 7.**
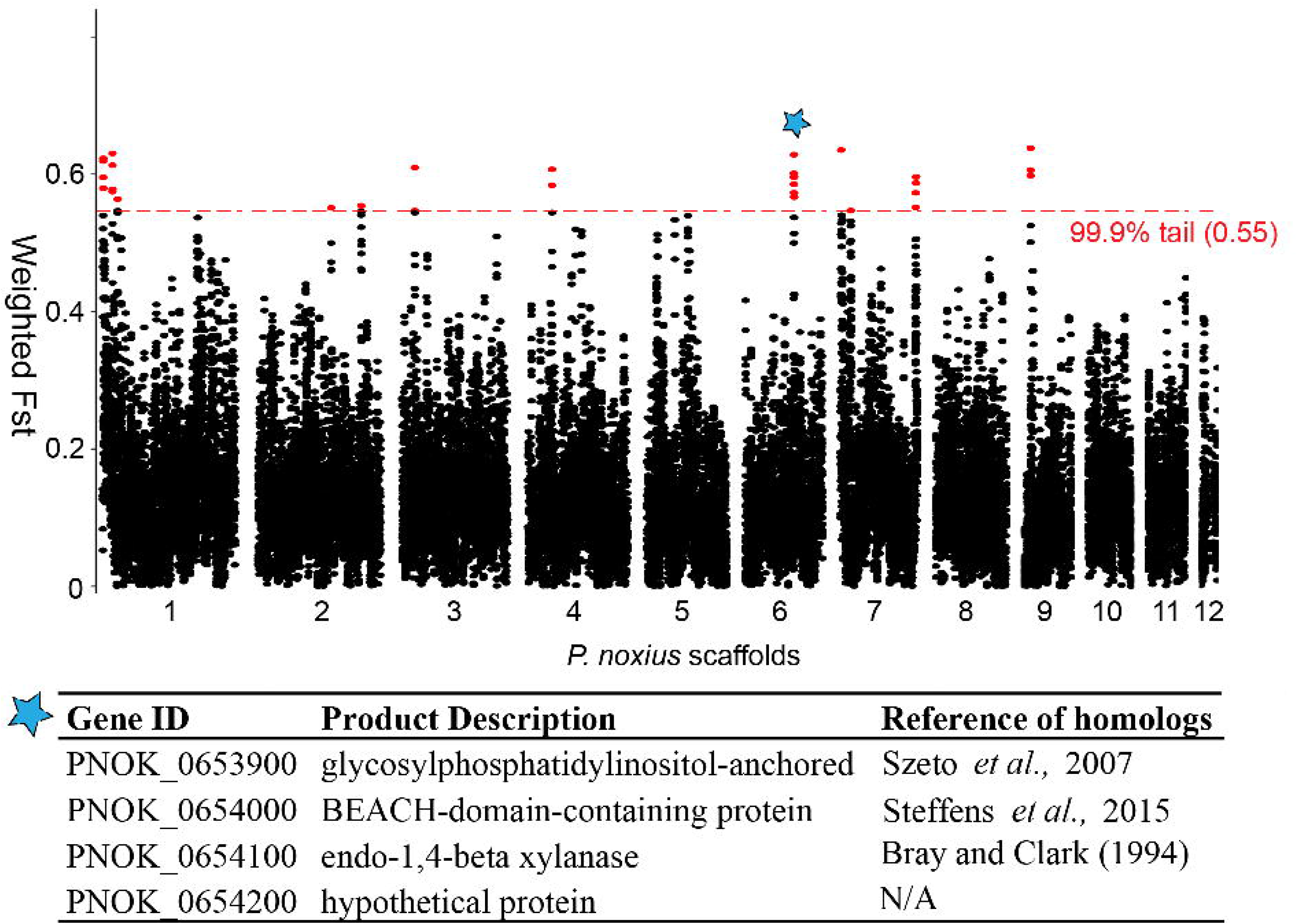
Weighted F_ST_ values for 5kb windows across the *P. noxius* assembly. Red colour dots indicate windows having F_ST_ value greater than 99.9% tail of 0.55. The 12-kb candidate region is marked in blue star. Annotations and references of genes located in this candidate region are listed.

## Discussion

Here we report four high-quality genome sequences of Hymenochaetales species that are global tree pathogens of particular importance. To date chromosome-level assemblies are available only for a few basidiomycetous species (Manuel Alfaro et al., 2016; Foulongne-Oriol et al., 2016; J. E. Stajich et al., 2010) including *P. noxius* (Table S5). Orthologous relationships with other complete genomes of basidiomycetes have confirmed conservation of karyotypes with few fusion or breaks in Agaricomycetes. Our study has shown the diversity and abundance of CAZymes, in particularly lignin-degrading enzymes, in the genomes of Hymenochaetaceae species. Such genetic architecture has been demonstrated in other white rot fungi (Riley et al., 2014), and differences in CAZymes have been implicated as the genetic basis of softwood or hardwood utilization (Suzuki et al., 2012). As revealed by comparative genomics analysis, *P. noxius* has highly expanded 1,3-beta-glucan synthase genes and abundant CAZymes, making it an aggressive wood decay pathogen of a wide variety of both broadleaved and coniferous trees. The results can serve as a starting point for understanding the ecological role of *P. noxius*. Our study also identified high copy numbers of C2-NACHT-WD40(n)-containing NLRs and candidate genes associated with population differentiation. The NLR family is highly diverse in fungi and was found to be central to the process of programmed cell death and implicated in fungal vegetative incompatibility and general nonself recognition (Bidard, Clave, & Saupe, 2013). Analysis of global transcription at different pathogenesis stages and detailed functional assays will help resolve their functions.

Population genomics analyses of *P. noxius* suggest that it is a hypervariable species. Our investigations into mating type loci and genome-wide heterozygosity further indicated that the genetic hyperdiversity can be attributed to a bipolar heterothallic reproductive system and heterokaryotic nature, though gene flow and/or high mutation rate may also play some role. The characteristic large interval between *HD* pairs has only been reported in *Schizophyllum commune* (~55 kb) and *Flammulina velutipes* (~70 kb), in which the genomic separation likely emerged through inversions or transpositions of gene clusters surrounding *HD* (van Peer et al., 2011). This exceptional large separation between *HD* pairs would allow higher probability of recombination between the physically distant *HD* genes(James et al., 2013; van Peer et al., 2011), thus resulting in progeny with more diverse mating types which are ready to mate. Notably, both monokaryotic and heterokaryotic state of *P. noxius* mycelia are prevalent in the nature, and some isolates likely contain more than two genetically different nuclei (Chung et al., 2015). In addition, some isolates were able to produce basidiocarps by themselves when cultured on sawdust medium (P.-J. Ann, pers. comm.), suggesting dikaryotization or homokaryotic fruiting occurred spontaneously or in response to certain conditions (Wendland, 2016). It would be interesting to further investigate the complex regulatory mechanisms underlying anastomosis, karyogamy, and meiotic division during vegetative growth and basidiocarp formation. Transcriptional diversity among genetically variable individuals is also warrant further exploration.

How *P. noxius* is spread in regions of Asia has been the focus of a few studies (Akiba et al., 2015; Chung et al., 2015; Hattori, Abe, & Usugi, 1996). Genomic analysis of the strains from across Taiwan and offshore islands of Japan allowed us to re-examine possible mode of disease transmission in the Asia Pacific region. The pattern of gene flow within and between islands suggested that human activity such as planting of infected seedlings may have promoted the movement of *P. noxius*. which provides an artificial environment for population to increase. The within-population hyperdiversity, prevalence of monokaryotic isolates, and sporadic pattern of new disease foci also support the involvement of basidiospores in *P. noxius* dissemination. Considering its host range and fast-growing ability in warm weather, and the abundant basidiospores that can be produced from perennial fruiting bodies with huge dispersal potential (e.g., the basidiospores of *Heterobasidion annosum* and *Peniophora aurantiaca* were captured 50–500 km and ~1000 km apart from the inoculum source), *P. noxius* may potentially affect more agricultural, ecological, and residential environments. A preliminary model based on 19 bioclimatic variables of known locations of ~100 *P. noxius* isolates from south eastern Asia, Australia, and Pacific islands predicted that extensive global regions are at risk, which includes a big part of the South American continent (Klopfenstein et al., 2016).

The Hymenochaetales is phylogenetically placed between the better-studied Agaricales and Ustillaginales orders, making the reference assembly of *P. noxius* KPN91 an attractive genome to study the evolutionary transition between these orders of Basidiomycota. It should be of continuous interest to confirm this observation when more reference genome assemblies become available. Genetic hyperdiversity of *P. noxius* suggests that the pathogen may have greater adaptability to different environments and stresses. Associating growth phenotypes under a variety of conditions to the variations identified in this study will be a logical next step for a full understanding of the genetic basis underlying the fitness and virulence of white root rot fungi. For disease control and prevention, much more attention needs to be paid to monitor how these fungi will behave in changing or extreme weather conditions.

## Availability of data and materials

Genome assemblies and annotations are available from NCBI under whole genome shotgun (WGS) ID: NBII00000000 (*P. noxius*), NBBA00000000 (*P. sulphurascens*), NBAY00000000 (*P. pini*) and NBAZ00000000 (*P. lamaensis*). Bioproject and Biosample ID of raw data are available in Table S1 and S2.

## Author’s contributions

Strain cultivation and preparation: H.H.L., T.J.L, H.M.K, M.A., T.H., Y.O., N.S. and T.K. Strain provider: C.L.C., R.F.L., S.S.T., P.J.A., J.N.T., M.A., T.H., Y.O., N.S. Strain sequencing and assembly: C.L.C., H.H.L., C.Y.C., M.J.L., T.K. and I.J.T. Annotation and manual curation: H.M.K., T.J.L and I.J.T. Comparative genomics analysis: T.H.K, D.L., M.B.R., H.M.K. and I.J.T. Population genomics analysis: H.M.K., T.J.L and I.J.T. Mating locus analysis: C.L.C., H.H.L., Y.Y.C., T.H.K., I.J.T. RNA-seq analysis: T.J.L and I.J.T. Wrote the manuscript: C.L.C, T.J.L, H.H.L, T.K and I.J.T. Conceived and directed the project: C.L.C., T.K. and I.J.T.

## Acknowledgements

We thank John Wang (Biodiversity Research Center, Academia Sinica), Yen-Ping Hsueh (IMB, Academia Sinica) and Robert Waterhouse (University of Lausanne) for commenting the manuscript. We thank Chian-mei Wei, Kuan-chun Chen (High Throughput Genomics Core at Biodiversity Research Center, Academia Sinica) for sequencing. We thank Tun-Tschu Chang and Yu-Ching Huang for strain collection and consultation.

## References

Akiba, M., Ota, Y., Tsai, I. J., Hattori, T., Sahashi, N., & Kikuchi, T. (2015). Genetic Differentiation and Spatial Structure of Phellinus noxius, the Causal Agent of Brown Root Rot of Woody Plants in Japan. Plos One, 10(10), e0141792. doi:10.1371/journal.pone.0141792

Alexa, A., & Rahnenfuhrer, J. (2016). topGO: Enrichment Analysis for Gene Ontology. R package version 2.26.0.

Alfaro, M., Castanera, R., Lavin, J. L., Grigoriev, I. V., Oguiza, J. A., Ramirez, L., & Pisabarro, A. G. (2016). Comparative and transcriptional analysis of the predicted secretome in the lignocellulose-degrading basidiomycete fungus Pleurotus ostreatus. Environ Microbiol, 18(12), 4710–4726. doi:10.1111/1462-2920.13360

Ann, P., Chang, T., & Ko, W. (2002). Phellinus noxius Brown Root Rot of Fruit and Ornamental Trees in Taiwan. Plant Disease, 86.

Ann, P. J., Lee, H. L., & Huang, T. C. (1999). Brown Root Rot of 10 Species of Fruit Trees Caused by Phellinus noxius in Taiwan. Plant Disease, 83(8), 746–750.

Apse, M. P., Aharon, G. S., Snedden, W. A., & Blumwald, E. (1999). Salt tolerance conferred by overexpression of a vacuolar Na+/H+ antiport in Arabidopsis. Science, 285(5431), 1256–1258.

Bankevich, A., Nurk, S., Antipov, D., Gurevich, A. A., Dvorkin, M., Kulikov, A. S., … Pevzner, P. A. (2012). SPAdes: A New Genome Assembly Algorithm and Its Applications to Single-Cell Sequencing. Journal of Computational Biology, 19(5), 455–477. doi:10.1089/cmb.2012.0021

Baranova, M. A., Logacheva, M. D., Penin, A. A., Seplyarskiy, V. B., Safonova, Y. Y., Naumenko, S. A., … Kondrashov, A. S. (2015). Extraordinary Genetic Diversity in a Wood Decay Mushroom. Mol Biol Evol, 32(10), 2775–2783. doi:10.1093/molbev/msv153

Bidard, F., Clave, C., & Saupe, S. J. (2013). The transcriptional response to nonself in the fungus Podospora anserina. G3 (Bethesda), 3(6), 1015–1030. doi:10.1534/g3.113.006262

Bolger, A. M., Lohse, M., & Usadel, B. (2014). Trimmomatic: a flexible trimmer for Illumina sequence data. Bioinformatics, 30(15), 2114–2120. doi:10.1093/bioinformatics/btu170

Borodovsky, M., & Lomsadze, A. (2011). Eukaryotic gene prediction using GeneMark.hmm-E and GeneMark-ES. Curr Protoc Bioinformatics, Chapter 4, Unit 4 6 1–10. doi:10.1002/0471250953.bi0406s35

Butler, J., MacCallum, I., Kleber, M., Shlyakhter, I. A., Belmonte, M. K., Lander, E. S., … Jaffe, D. B. (2008). ALLPATHS: de novo assembly of whole-genome shotgun microreads. Genome Res, 18(5), 810–820. doi:10.1101/gr.7337908

Castanera, R., Lopez-Varas, L., Borgognone, A., LaButti, K., Lapidus, A., Schmutz, J., … Ramirez, L. (2016). Transposable Elements versus the Fungal Genome: Impact on Whole-Genome Architecture and Transcriptional Profiles. PLoS Genet, 12(6), e1006108. doi:10.1371/journal.pgen.1006108

Chain, P. S., Grafham, D. V., Fulton, R. S., Fitzgerald, M. G., Hostetler, J., Muzny, D., … Detter, J. C. (2009). Genomics. Genome project standards in a new era of sequencing. Science, 326(5950), 236–237. doi:10.1126/science.1180614

Chang, C. C., Chow, C. C., Tellier, L. C., Vattikuti, S., Purcell, S. M., & Lee, J. J. (2015). Second-generation PLINK: rising to the challenge of larger and richer datasets. Gigascience, 4, 7. doi:10.1186/s13742-015-0047-8

Chang, T.-T. (1992). Decline of some forest trees associated with brown root rot caused by Phellinus noxius. Plant Pathology Bulletin, 1(2), 90–95.

Chin, C.-S., Alexander, D. H., Marks, P., Klammer, A. A., Drake, J., Heiner, C., … Korlach, J. (2013). Nonhybrid, finished microbial genome assemblies from long-read SMRT sequencing data. Nature Methods, 10(6), 563–569. doi:10.1038/nmeth.2474

Chin, C.-S., Peluso, P., Sedlazeck, F. J., Nattestad, M., Concepcion, G. T., Clum, A., … Schatz, M. C. (2016). Phased diploid genome assembly with single-molecule real-time sequencing. Nature Methods, 13(12), 1050–1054. doi:10.1038/nmeth.4035

Chung, C.-L., Huang, S.-Y., Huang, Y.-C., Tzean, S.-S., Ann, P.-J., Tsai, J.-N., … Liou, R.-F. (2015). The Genetic Structure of Phellinus noxius and Dissemination Pattern of Brown Root Rot Disease in Taiwan. Plos One, 10(10), e0139445. doi:10.1371/journal.pone.0139445

Conesa, A., Gotz, S., Garcia-Gomez, J. M., Terol, J., Talon, M., & Robles, M. (2005). Blast2GO: a universal tool for annotation, visualization and analysis in functional genomics research. Bioinformatics, 21(18), 3674–3676. doi:10.1093/bioinformatics/bti610

Cunniffe, N. J., Cobb, R. C., Meentemeyer, R. K., Rizzo, D. M., & Gilligan, C. A. (2016). Modeling when, where, and how to manage a forest epidemic, motivated by sudden oak death in California. Proc Natl Acad Sci U S A, 113(20), 5640–5645. doi:10.1073/pnas.1602153113

Danecek, P., Auton, A., Abecasis, G., Albers, C. A., Banks, E., DePristo, M. A., … Genomes Project Analysis, G. (2011). The variant call format and VCFtools. Bioinformatics, 27(15), 2156–2158. doi:10.1093/bioinformatics/btr330

Dashtban, M., Schraft, H., Syed, T. A., & Qin, W. (2010). Fungal biodegradation and enzymatic modification of lignin. Int J Biochem Mol Biol, 1(1), 36–50.

Douglas, C. M. (2001). Fungal beta(1,3)-D-glucan synthesis. Med Mycol, 39 Suppl 1, 55–66.

Emms, D. M., & Kelly, S. (2015). OrthoFinder: solving fundamental biases in whole genome comparisons dramatically improves orthogroup inference accuracy. Genome Biol, 16, 157. doi:10.1186/s13059-015-0721-2

Felsenstein, J. (2005). PHYLIP (Phylogeny Inference Package) version 3.6. Distributed by the author. Department of Genome Sciences, University of Washington, Seattle.

Finn, R. D., Coggill, P., Eberhardt, R. Y., Eddy, S. R., Mistry, J., Mitchell, A. L., … Bateman, A. (2016). The Pfam protein families database: towards a more sustainable future. Nucleic Acids Research, 44(D1), D279–D285. doi:10.1093/nar/gkv1344

Fisher, M. C., Henk, D. A., Briggs, C. J., Brownstein, J. S., Madoff, L. C., McCraw, S. L., & Gurr, S. J. (2012). Emerging fungal threats to animal, plant and ecosystem health. Nature, 484(7393), 186–194. doi:10.1038/nature10947

Floudas, D., Binder, M., Riley, R., Barry, K., Blanchette, R. A., Henrissat, B., … Hibbett, D. S. (2012). The Paleozoic origin of enzymatic lignin decomposition reconstructed from 31 fungal genomes. Science, 336(6089), 1715–1719. doi:10.1126/science.1221748

Foulongne-Oriol, M., Rocha de Brito, M., Cabannes, D., Clément, A., Spataro, C., Moinard, M., … Savoie, J.-M. (2016). The Genetic Linkage Map of the Medicinal Mushroom Agaricus subrufescens Reveals Highly Conserved Macrosynteny with the Congeneric Species Agaricus bisporus. G3: Genes|Genomes|Genetics, 6(5), 1217–1226. doi:10.1534/g3.115.025718

Garrison, E., & Marth, G. (2012). Haplotype-based variant detection from short-read sequencing. arXiv, 1207, 3907.

Goberville, E., Hautekeete, N. C., Kirby, R. R., Piquot, Y., Luczak, C., & Beaugrand, G. (2016). Climate change and the ash dieback crisis. Sci Rep, 6, 35303. doi:10.1038/srep35303

Grigoriev, I. V., Nikitin, R., Haridas, S., Kuo, A., Ohm, R., Otillar, R., … Shabalov, I. (2014). MycoCosm portal: gearing up for 1000 fungal genomes. Nucleic Acids Research, 42(D1), D699–D704. doi:10.1093/nar/gkt1183

Gross, A., Holdenrieder, O., Pautasso, M., Queloz, V., & Sieber, T. N. (2014). Hymenoscyphus pseudoalbidus, the causal agent of European ash dieback. Mol Plant Pathol, 15(1), 5–21. doi:10.1111/mpp.12073

Guy, L., Roat Kultima, J., & Andersson, S. G. E. (2010). genoPlotR: comparative gene and genome visualization in R. Bioinformatics, 26(18), 2334–2335. doi:10.1093/bioinformatics/btq413

Hane, J. K., Rouxel, T., Howlett, B. J., Kema, G. H., Goodwin, S. B., & Oliver, R. P. (2011). A novel mode of chromosomal evolution peculiar to filamentous Ascomycete fungi. Genome Biol, 12(5), R45. doi:10.1186/gb-2011-12-5-r45

Hattori, T., Abe, Y., & Usugi, T. (1996). Distribution of clones of Phellinus noxius in a windbreak on Ishigaki Island. European Journal of Forest Pathology, 26(2), 69–80. doi:10.1111/j.1439-0329.1996.tb00711.x

Hepting, G. H. (1971). Diseases of forest and shade trees of the United States: U.S. Dept. Of Agriculture Forest Service Handbook Number 386.

Hoff, K. J., Lange, S., Lomsadze, A., Borodovsky, M., & Stanke, M. (2016). BRAKER1: Unsupervised RNA-Seq-Based Genome Annotation with GeneMark-ET and AUGUSTUS. Bioinformatics, 32(5), 767–769. doi:10.1093/bioinformatics/btv661

Huang, H., Sun, L., Bi, K., Zhong, G., & Hu, M. (2016). The effect of phenazine-1-carboxylic acid on the morphological, physiological, and molecular characteristics of Phellinus noxius. Molecules, 21(5), 613.

Hunt, M., Kikuchi, T., Sanders, M., Newbold, C., Berriman, M., & Otto, T. D. (2013). REAPR: a universal tool for genome assembly evaluation. Genome biology, 14, R47. doi:10.1186/gb-2013-14-5-r47

James, T. Y., Lee, M., & van Diepen, L. T. (2011). A single mating-type locus composed of homeodomain genes promotes nuclear migration and heterokaryosis in the white-rot fungus Phanerochaete chrysosporium. Eukaryot Cell, 10(2), 249–261. doi:10.1128/EC.00212-10

James, T. Y., Srivilai, P., Kues, U., & Vilgalys, R. (2006). Evolution of the bipolar mating system of the mushroom Coprinellus disseminatus from its tetrapolar ancestors involves loss of mating-type-specific pheromone receptor function. Genetics, 172(3), 1877–1891. doi:10.1534/genetics.105.051128

James, T. Y., Sun, S., Li, W., Heitman, J., Kuo, H. C., Lee, Y. H., … Olson, A. (2013). Polyporales genomes reveal the genetic architecture underlying tetrapolar and bipolar mating systems. Mycologia, 105(6), 1374–1390. doi:10.3852/13-162

Jancova, P., Anzenbacher, P., & Anzenbacherova, E. (2010). Phase II drug metabolizing enzymes. Biomed Pap Med Fac Univ Palacky Olomouc Czech Repub, 154(2), 103–116.

Jones, P., Binns, D., Chang, H.-Y., Fraser, M., Li, W., McAnulla, C., … Hunter, S. (2014). InterProScan 5: genome-scale protein function classification. Bioinformatics, 30(9), 1236–1240. doi:10.1093/bioinformatics/btu031

Kampe, J., Kahmann, R., Bolker, M., Ma, L. J., Brefort, T., Saville, B. J., … Birren, B. W. (2006). Insights from the genome of the biotrophic fungal plant pathogen Ustilago maydis. Nature, 444(7115), 97–101. doi:10.1038/nature05248

Katoh, K., & Standley, D. M. (2014). MAFFT: iterative refinement and additional methods. Methods Mol Biol, 1079, 131–146. doi:10.1007/978-1-62703-646-7_8

Klopfenstein, N. B., Pitman, E. W. I., Hanna, J. W., Cannon, P. G., Stewart, J. E., Sahashi, N., … Kim, M.-S. (2016). A preliminary bioclimatic approach to predicting potential distribution of Phellinus noxious and geographical areas at risk from invasion. Paper presented at the Proceedings of the 63rd Annual Western International Forest Disease Work Conference.

Koboldt, D. C., Zhang, Q., Larson, D. E., Shen, D., McLellan, M. D., Lin, L., … Wilson, R. K. (2012). VarScan 2: Somatic mutation and copy number alteration discovery in cancer by exome sequencing. Genome research, 22(3), 568–576. doi:10.1101/gr.129684.111

Kües, U. (2015). From two to many: multiple mating types in Basidiomycetes. Fungal Biology Reviews, 29(3–4), 126–166. doi:http://dx.doi.org/10.1016/j.fbr.2015.11.001

Kumar, S., Stecher, G., & Tamura, K. (2016). MEGA7: Molecular Evolutionary Genetics Analysis Version 7.0 for Bigger Datasets. Mol Biol Evol, 33(7), 1870–1874. doi:10.1093/molbev/msw054

Kurtz, S., Phillippy, A., Delcher, A. L., Smoot, M., Shumway, M., Antonescu, C., & Salzberg, S. L. (2004). Versatile and open software for comparing large genomes. Genome Biology, 5(2), R12.

Lam, K. K., LaButti, K., Khalak, A., & Tse, D. (2015). FinisherSC: a repeat-aware tool for upgrading de novo assembly using long reads. Bioinformatics, 31(19), 3207–3209. doi:10.1093/bioinformatics/btv280

Larsson, K.-H., Parmasto, E., Fischer, M., Langer, E., Nakasone, K. K., & Redhead, S. a. (2006). Hymenochaetales: a molecular phylogeny for the hymenochaetoid clade. Mycologia, 98, 926–936.

Lavezzo, E., Falda, M., Fontana, P., Bianco, L., & Toppo, S. (2016). Enhancing protein function prediction with taxonomic constraints–The Argot2.5 web server. Methods, 93, 15–23. doi:10.1016/j.ymeth.2015.08.021

Leffler, E. M., Bullaughey, K., Matute, D. R., Meyer, W. K., Segurel, L., Venkat, A., … Przeworski, M. (2012). Revisiting an old riddle: what determines genetic diversity levels within species? PLoS Biol, 10(9), e1001388. doi:10.1371/journal.pbio.1001388

Leipe, D. D., Koonin, E. V., & Aravind, L. (2004). STAND, a class of P-loop NTPases including animal and plant regulators of programmed cell death: multiple, complex domain architectures, unusual phyletic patterns, and evolution by horizontal gene transfer. J Mol Biol, 343(1), 1–28. doi:10.1016/j.jmb.2004.08.023

Li, H., Handsaker, B., Wysoker, A., Fennell, T., Ruan, J., Homer, N., … Durbin, R. (2009). The Sequence Alignment/Map format and SAMtools. Bioinformatics (Oxford, England), 25, 2078–2079. doi:10.1093/bioinformatics/btp352

Loftus, B. J., Fung, E., Roncaglia, P., Rowley, D., Amedeo, P., Bruno, D., … Hyman, R. W. (2005). The genome of the basidiomycetous yeast and human pathogen Cryptococcus neoformans. Science, 307(5713), 1321–1324. doi:10.1126/science.1103773

Martin, F., Aerts, A., Ahren, D., Brun, A., Danchin, E. G., Duchaussoy, F., … Grigoriev, I. V. (2008). The genome of Laccaria bicolor provides insights into mycorrhizal symbiosis. Nature, 452(7183), 88–92. doi:10.1038/nature06556

Maurer-Stroh, S., & Eisenhaber, F. (2005). Refinement and prediction of protein prenylation motifs. Genome biology, 6(6), R55.

Min, B., Park, H., Jang, Y., Kim, J. J., Kim, K. H., Pangilinan, J., … Choi, I. G. (2015). Genome sequence of a white rot fungus Schizopora paradoxa KUC8140 for wood decay and mycoremediation. J Biotechnol, 211, 42–43. doi:10.1016/j.jbiotec.2015.06.426

Morin, E., Kohler, A., Baker, A. R., Foulongne-Oriol, M., Lombard, V., Nagy, L. G., … Martin, F. (2012). Genome sequence of the button mushroom Agaricus bisporus reveals mechanisms governing adaptation to a humic-rich ecological niche. Proc Natl Acad Sci U S A, 109(43), 17501–17506. doi:10.1073/pnas.1206847109

Nagy, L. G., Riley, R., Tritt, A., Adam, C., Daum, C., Floudas, D., … Hibbett, D. S. (2016). Comparative Genomics of Early-Diverging Mushroom-Forming Fungi Provides Insights into the Origins of Lignocellulose Decay Capabilities. Mol Biol Evol, 33(4), 959–970. doi:10.1093/molbev/msv337

Nandris, D., Nicole, M., & Geiger, J. P. (1987). Variation in virulence among Rigidoporus lignosus and Phellinus noxius isolates from West Africa1. European Journal of Forest Pathology, 17(4-5), 271–281. doi:10.1111/j.1439-0329.1987.tb01026.x

Niculita-Hirzel, H., Labbe, J., Kohler, A., le Tacon, F., Martin, F., Sanders, I. R., & Kues, U. (2008). Gene organization of the mating type regions in the ectomycorrhizal fungus Laccaria bicolor reveals distinct evolution between the two mating type loci. New Phytol, 180(2), 329–342. doi:10.1111/j.1469-8137.2008.02525.x

Ohm, R. A., de Jong, J. F., Lugones, L. G., Aerts, A., Kothe, E., Stajich, J. E., … Wosten, H. A. (2010). Genome sequence of the model mushroom Schizophyllum commune. Nat Biotechnol, 28(9), 957–963. doi:10.1038/nbt.1643

Olson, A., Aerts, A., Asiegbu, F., Belbahri, L., Bouzid, O., Broberg, A., … Stenlid, J. (2012). Insight into trade-off between wood decay and parasitism from the genome of a fungal forest pathogen. New Phytol, 194(4), 1001–1013. doi:10.1111/j.1469-8137.2012.04128.x

Parra, G., Bradnam, K., & Korf, I. (2007). CEGMA: a pipeline to accurately annotate core genes in eukaryotic genomes. Bioinformatics (Oxford, England), 23, 1061–1067. doi:10.1093/bioinformatics/btm071

Potter, C., Harwood, T., Knight, J., & Tomlinson, I. (2011). Learning from history, predicting the future: the UK Dutch elm disease outbreak in relation to contemporary tree disease threats. Philos Trans R Soc Lond B Biol Sci, 366(1573), 1966–1974. doi:10.1098/rstb.2010.0395

Price, M. N., Dehal, P. S., & Arkin, A. P. (2009). FastTree: Computing Large Minimum Evolution Trees with Profiles instead of a Distance Matrix. Molecular Biology and Evolution, 26(7), 1641–1650. doi:10.1093/molbev/msp077

Raj, A., Stephens, M., & Pritchard, J. K. (2014). fastSTRUCTURE: Variational Inference of Population Structure in Large SNP Data Sets. Genetics, 197(2), 573–589. doi:10.1534/genetics.114.164350

Riley, R., Salamov, a. a., Brown, D. W., Nagy, L. G., Floudas, D., Held, B. W., … Grigoriev, I. V. (2014b). Extensive sampling of basidiomycete genomes demonstrates inadequacy of the white-rot/brown-rot paradigm for wood decay fungi. Proceedings of the National Academy of Sciences. doi:10.1073/pnas.1400592111

Sahashi, N., Akiba, M., Ishihara, M., Miyazaki, K., & Kanzaki, N. (2010). Cross Inoculation Tests with Phellinus noxius Isolates from Nine Different Host Plants in the Ryukyu Islands, Southwestern Japan. Plant Disease, 94(3), 358–360.

Sahashi, N., Akiba, M., Ishihara, M., Ota, Y., & Kanzaki, N. (2012). Brown root rot of trees caused by Phellinus noxius in the Ryukyu Islands, subtropical areas of Japan. Forest Pathology, 42, 353–361.

Sahashi, N., Akiba, M., Takemoto, S., Yokoi, T., Ota, Y., & Kanzaki, N. (2014). Phellinus noxius causes brown root rot on four important conifer species in Japan. European Journal of Plant Pathology, 140(4), 869–873.

Schwartz, S., Zhang, Z., Frazer, K. A., Smit, A., Riemer, C., Bouck, J., … Miller, W. (2000). PipMaker–a web server for aligning two genomic DNA sequences. Genome Research, 10(4), 577–586.

Schwarze, F., Jauss, F., Spencer, C., Hallam, C., & Schubert, M. (2012). Evaluation of an antagonistic Trichoderma strain for reducing the rate of wood decomposition by the white rot fungus Phellinus noxius. Biological Control, 61(2), 160–168.

Simão, F. A., Waterhouse, R. M., Ioannidis, P., Kriventseva, E. V., & Zdobnov, E. M. (2015). BUSCO: assessing genome assembly and annotation completeness with single-copy orthologs. Bioinformatics, 31(19), 3210–3212. doi:10.1093/bioinformatics/btv351

Stajich, J. E., Wilke, S. K., Ahren, D., Au, C. H., Birren, B. W., Borodovsky, M., … Pukkila, P. J. (2010). Insights into evolution of multicellular fungi from the assembled chromosomes of the mushroom Coprinopsis cinerea (Coprinus cinereus). Proceedings of the National Academy of Sciences, 107(26), 11889–11894. doi:10.1073/pnas.1003391107

Stamatakis, A. (2006). RAxML-VI-HPC: maximum likelihood-based phylogenetic analyses with thousands of taxa and mixed models. Bioinformatics (Oxford, England), 22, 2688–2690. doi:10.1093/bioinformatics/btl446

Stanke, M., Tzvetkova, A., & Morgenstern, B. (2006). AUGUSTUS at EGASP: using EST, protein and genomic alignments for improved gene prediction in the human genome. Genome biology, 7 Suppl 1, S11.11–18. doi:10.1186/gb-2006-7-s1-s11

Steffens, A., Brautigam, A., Jakoby, M., & Hulskamp, M. (2015). The BEACH Domain Protein SPIRRIG Is Essential for Arabidopsis Salt Stress Tolerance and Functions as a Regulator of Transcript Stabilization and Localization. PLoS Biol, 13(7), e1002188. doi:10.1371/journal.pbio.1002188

Suzuki, H., MacDonald, J., Syed, K., Salamov, A., Hori, C., Aerts, A., … Master, E. R. (2012). Comparative genomics of the white-rot fungi, Phanerochaete carnosa and P. chrysosporium, to elucidate the genetic basis of the distinct wood types they colonize. BMC Genomics, 13(1). doi:10.1186/1471-2164-13-444

Szeto, C. Y., Leung, G. S., & Kwan, H. S. (2007). Le.MAPK and its interacting partner, Le.DRMIP, in fruiting body development in Lentinula edodes. Gene, 393(1-2), 87–93. doi:10.1016/j.gene.2007.01.030

Van der Nest, M. A., Olson, A., Lind, M., Velez, H., Dalman, K., Brandstrom Durling, M., … Stenlid, J. (2014). Distribution and evolution of het gene homologs in the basidiomycota. Fungal Genet Biol, 64, 45–57. doi:10.1016/j.fgb.2013.12.007

van Peer, A. F., Park, S.-Y., Shin, P.-G., Jang, K.-Y., Yoo, Y.-B., Park, Y.-J., … Kong, W.-S. (2011). Comparative genomics of the mating-type loci of the mushroom Flammulina velutipes reveals widespread synteny and recent inversions. Plos One, 6(7), e22249. doi:10.1371/journal.pone.0022249

Vilella, A. J., Blanco-Garcia, A., Hutter, S., & Rozas, J. (2005). VariScan: Analysis of evolutionary patterns from large-scale DNA sequence polymorphism data. Bioinformatics, 21(11), 2791–2793. doi:10.1093/bioinformatics/bti403

Walker, B. J., Abeel, T., Shea, T., Priest, M., Abouelliel, A., Sakthikumar, S., … Earl, A. M. (2014). Pilon: An Integrated Tool for Comprehensive Microbial Variant Detection and Genome Assembly Improvement. Plos One, 9(11), e112963. doi:10.1371/journal.pone.0112963

Weber, T., Blin, K., Duddela, S., Krug, D., Kim, H. U., Bruccoleri, R., … Medema, M. H. (2015). antiSMASH 3.0—a comprehensive resource for the genome mining of biosynthetic gene clusters. Nucleic Acids Research, 43(W1), W237–W243. doi:10.1093/nar/gkv437

Wences, A. H., & Schatz, M. C. (2015). Metassembler: merging and optimizing de novo genome assemblies. Genome biology, 16(1). doi:10.1186/s13059-015-0764-4

Wendland, J. (2016). Growth, Differentiation and Sexuality (J. Wendland Ed. 3 ed.): Springer.

Williams, H. L., Sturrock, R. N., Islam, M. A., Hammett, C., Ekramoddoullah, A. K., & Leal, I. (2014). Gene expression profiling of candidate virulence factors in the laminated root rot pathogen Phellinus sulphurascens. BMC Genomics, 15, 603. doi:10.1186/1471-2164-15-603

Wu, J., Peng, S. L., Zhao, H. B., Tang, M. H., Li, F. R., & Chen, B. M. (2011). Selection of species resistant to the wood rot fungus Phellinus noxius. European Journal of Plant Pathology, 130(4), 463–467. doi:10.1007/s10658-011-9774-6

Yin, Y., Mao, X., Yang, J., Chen, X., Mao, F., & Xu, Y. (2012). dbCAN: a web resource for automated carbohydrate-active enzyme annotation. Nucleic Acids Res, 40(Web Server issue), W445–451. doi:10.1093/nar/gks479

Zhou, L.-W., Vlasák, J., & Dai, Y.-C. (2016). Taxonomy and phylogeny of Phellinidium (Hymenochaetales, Basidiomycota): A redefinition and the segregation of Coniferiporia gen. nov. for forest pathogens. Fungal Biology, 120(8), 988–1001. doi:http://dx.doi.org/10.1016/j.funbio.2016.04.008

Zuccaro, A., Lahrmann, U., Guldener, U., Langen, G., Pfiffi, S., Biedenkopf, D., … Kogel, K. H. (2011). Endophytic life strategies decoded by genome and transcriptome analyses of the mutualistic root symbiont Piriformospora indica. PLoS Pathog, 7(10), e1002290. doi:10.1371/journal.ppat.1002290

